# Widespread *cis*-regulation of RNA-editing in a large mammal

**DOI:** 10.1101/304220

**Authors:** Thomas J Lopdell, Christine Couldrey, Kathryn Tiplady, Stephan R Davis, Russell G Snell, Bevin L Harris, Mathew D Littlejohn

**Affiliations:** Research & Development, Livestock Improvement Corporation, Hamilton, New Zealand; School of Biological Sciences, University of Auckland, Auckland, New Zealand

**Keywords:** RNA editing, GWAS, Milk, QTL mapping, RNA sequencing, Genome sequencing

## Abstract

Post-transcriptional RNA editing may regulate transcript expression and diversity in cells, with potential impacts on various aspects of physiology and environmental adaptation. A small number of recent genome-wide studies in *Drosophila*, mouse, and human have shown that RNA editing can be genetically modulated, highlighting loci that quantitatively impact editing of transcripts. The potential gene expression and physiological consequences of these RNA editing quantitative trait loci (edQTL), however, are almost entirely unknown. Here, we present analyses of RNA editing in a large domestic mammal *(Bos taurus)*, where we use whole genome and high depth RNA sequencing to discover, characterise, and conduct genetic mapping studies of novel transcript edits. Using a discovery population of nine deeply-sequenced cows, we identify 2,001 edit sites in the mammary transcriptome, the majority of which are adenosine to inosine edits (97.4%). Most sites are predicted to reside in double-stranded secondary structures (85.7%), and quantification of the rates of editing in an additional 355 cows reveals editing is negatively correlated with gene expression in the majority of cases. Genetic analyses of RNA editing and gene expression highlights 67 *cis*-regulated edQTL, of which seven appear to co-segregate with expression QTL effects. Trait association analyses in a separate population of 9,988 lactating cows also shows nine of the *cis*-edQTL coincide with at least one co-segregating lactation QTL. Together, these results enhance our understanding of RNA editing dynamics in mammals, and suggest mechanistic links by which loci may impact phenotype through RNA-editing mediated processes.

## Introduction

The process of gene expression involves transcribing the information stored in DNA into messenger RNA (mRNA). In Eukaryotes, most mRNA sequences differ to those of DNA, primarily due to RNA splicing. However, the process of RNA editing can add additional diversity, whereby bases in the transcript are altered *in-situ* by direct enzymatic modification. In metazoan cells, the most common form of RNA editing is deamination of adenosine (A) nucleotides, forming inosine (I), catalysed by enzymes from the adenosine deaminase acting on RNA *(ADAR)* family (Savva et al., 2012).

Depending on the location of edits within the pre-mRNA transcript, the potential consequences of RNA editing can include changes to the coding sequence, the creation or destruction of splice sites (Nishikura, 2010), triggering of nuclear retention mechanisms of edited transcripts (Zhang and Carmichael, 2001; Prasanth et al., 2005), or the creation or destruction of miRNA binding sites within the 3’-UTR (Liang and Landweber, 2007; Wang et al., 2013). These changes in turn can affect gene expression, either as part of normal regulation (Goldstein et al., 2017), in a pathogenic context such as cancer (Zhang et al., 2016; Baysal et al., 2017), or as a mechanism to regulate alternative splicing (Solomon et al., 2013).

Genetic regulation of gene expression, whether operating through polymorphic variation in *cis* or *trans* regulatory elements, or through other mechanisms such as DNA methylation, is thought to account for the majority of genetic variance in phenotypic traits (Nicolae et al., 2010). Identification of expression quantitative trait loci (eQTL), therefore, provides insight into causative mechanisms for co-locating QTL for more broadly defined physiological traits, where these methods have been applied to identify causative genes for various characters and diseases in humans (Zhu et al., 2016; Li et al., 2015; Li and Huang, 2017), model species (Parks et al., 2013, 2015), and agricultural and domestic species (Li et al., 2013; Littlejohn et al., 2016; Lopdell et al., 2017).

Since numerous regulatory effects have been attributed to RNA editing, genetic regulation of editing poses another potential mechanism to explain impacts on physiological traits. In three recent studies conducted in *Drosophila* (Ramaswami et al., 2015), mice (Gu et al., 2016), and humans (Park et al., 2017), researchers demonstrated the application of QTL mapping approaches to reveal widespread genetic modulation of RNA-editing. In the current study, we aimed to build on these studies by characterising the genetic landscape of RNA-editing in cattle, and more specifically, use these data to investigate potential regulatory effects of identified loci on gene expression and complex quantitative traits. Utilising whole genome sequencing, high depth mammary RNA sequencing, and genome-wide association approaches in outbred cattle populations, we report the *de novo* discovery of RNA edits, RNA editing QTL (edQTL), and a number of co-locating, co-segregating gene expression and lactation impacts as potential consequences of these modifications.

## Results

### Discovery and molecular context of edited sites

To identify candidate RNA editing sites, we performed whole genome sequencing of nine animals for which high depth RNAseq data was also available. Animals were sequenced at an average 22-fold read depth for genomic sequence and 104 million read pairs for RNAseq, with variants called for both DNA and RNA sequence alignments (Materials and Methods). Variants that were identified from RNA data but found to be absent from DNA data for the same animal were considered candidate sites. After applying further quality filtering to the variants (including visual inspection of alignments, see Materials and Methods) a total of 2,001 edited sites were identified. Edits mapped to a total of 314 genes (median 4.0 sites per gene, mean 6.4) with the majority of sites (85.1%) contained in intronic sequences (Table 1). Edits locating to the 3’ UTR were the next most common class (11.2%), with comparatively few edited sites in the 5’ UTR or coding exons. Relatively few sites were predicted to impact protein sequences (25 missense, 30 synonymous). These distributions are in broad agreement with previous reports of the distribution of edits in the human transcriptome (Chen, 2013).

**Table 1.** Locations of unambiguous edit sites.

| 5′UTR | Intron | Synonymous | Missense | 3′UTR |
| --- | --- | --- | --- | --- |
| 13 | 1561 | 30 | 25 | 205 |

Variant Effect Predictor results for 1,834 edited sites with unambiguous results, i.e., excluding those with alternative transcripts where multiple effects were predicted, and those where no gene on the correct strand was assigned. Variants for which the predictions were `upstream_gene_variant` or `downstream_gene_variant` were included in the 5′ or 3′ UTR respectively, due to imprecise annotation of these features for most bovine genes.

Of the different classes of base substitutions, A-to-I edits were by far the most common, (97.4% of sites; Table 2). Interestingly, however, the A-to-I edit class was much less dominant when only exonic sites are considered (32.7%, similar to the 40.7% reported elsewhere for a much larger sample (Chen, 2013)). In fact, 37 of the 52 non-A-to-I edited sites identified were exonic, raising the possibility that reads containing these edits arise from the expression of near-duplicate genes or pseudogenes, and have been incorrectly mapped. The most prevalent non A-to-I edits were G-to-A and C-to-U, concordant with previous literature for both humans (Chen, 2013) and cattle (Chen et al., 2016). As A-to-I edits are catalysed by the ADAR1 and ADAR2 enzymes, we confirmed the expression of the corresponding genes in mammary tissue, where the *ADAR1* gene was approximately 1.6-fold more highly expressed than *ADAR2* (Table 3). Minimal levels of expression were observed for homologues to human *APOBEC* genes, which have been implicated in non-A-to-I edits.

**Table 2.**
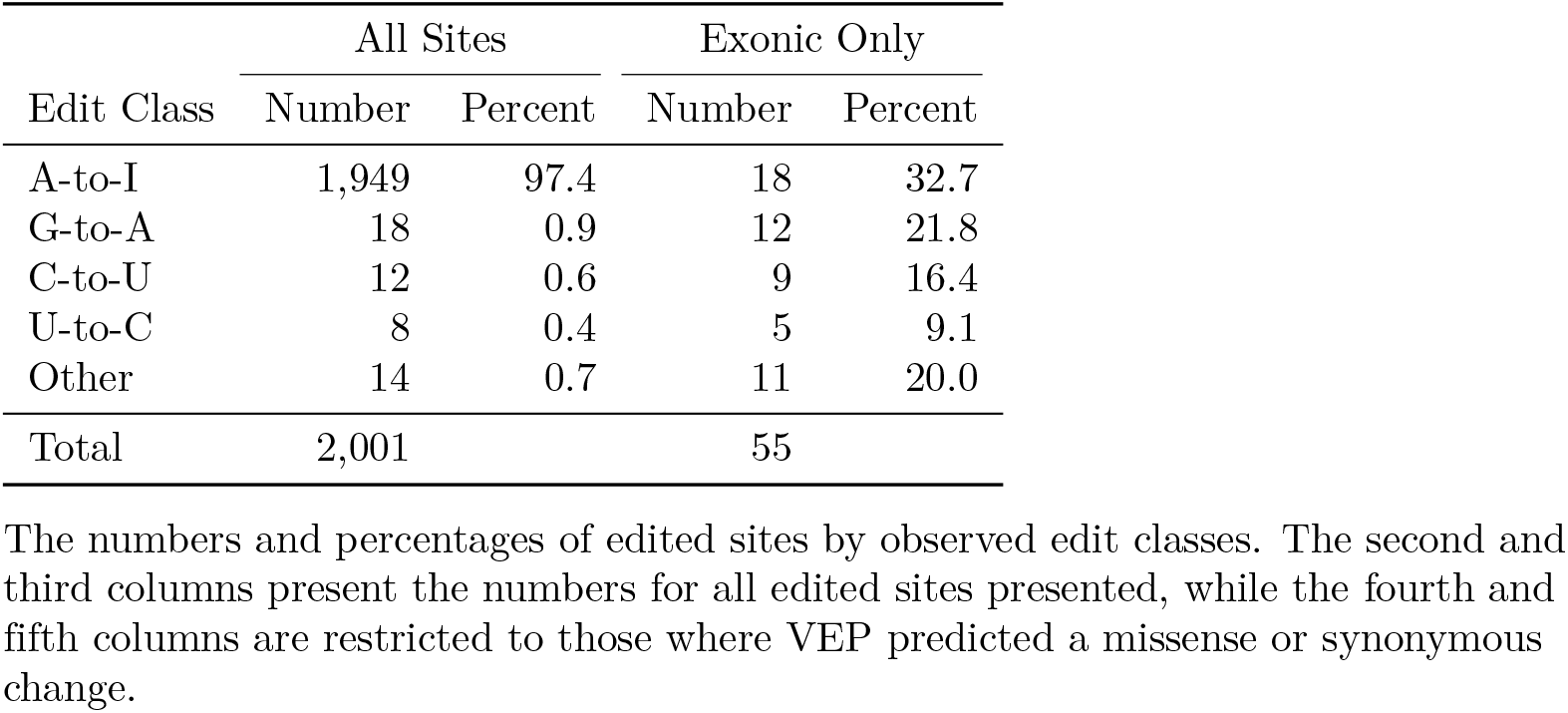
Observed numbers and frequencies of RNA edits by base.

| Edit Class | All Sites |  | Exonic Only |  |
| --- | --- | --- | --- | --- |
|  | Number | Percent | Number | Percent |
| A-to-I | 1,949 | 97.4 | 18 | 32.7 |
| G-to-A | 18 | 0.9 | 12 | 21.8 |
| C-to-U | 12 | 0.6 | 9 | 16.4 |
| U-to-C | 8 | 0.4 | 5 | 9.1 |
| Other | 14 | 0.7 | 11 | 20.0 |
| Total | 2,001 |  | 55 |  |

The numbers and percentages of edited sites by observed edit classes. The second and third columns present the numbers for all edited sites presented, while the fourth and fifth columns are restricted to those where VEP predicted a missense or synonymous change.

**Table 3.** Expression of ADAR genes in bovine mammary RNAseq.

| Gene | FPKM | TPM |
| --- | --- | --- |
| ADAR1 (ADAR) | $3.311 \pm 0.054$ | $4.081 \pm 0.082$ |
| ADAR2 (ADARB1) | $2.300 \pm 0.041$ | $2.824 \pm 0.057$ |
| ADAR3 (ADARB2) | $0.013 \pm 0.001$ | $0.016 \pm 0.001$ |
| APOBEC3A | $0.007 \pm 0.001$ | $0.009 \pm 0.001$ |
| APOBEC1 | $0.001 \pm 0.000$ | $0.001 \pm 0.001$ |

FPKM and TPM measures of gene expression in the lactating bovine mammary gland, for the 355 quantification animals. Values are shown as mean ± SEM. Gene names are given using both the traditional and HGNC (in brackets) nomenclatures. Expression levels are also displayed for the homologues of two human APOBEC genes that have been implicated in non-A-to-I editing: APOBEC3A (homologue ENSBTAG00000037800) in G-to-A editing, and APOBEC1 (homologue ENSBTAG00000014683) in C-to-U editing.

Non-uniform base usage was seen for bases directly adjacent to RNA editing sites (Table 4). In particular, guanosine was significantly under-represented at the position immediately upstream from edit sites (5.8% of bases; *p =* 3.63 × 10^−87^), but significantly enriched at positions immediately downstream (49.3% of bases; *p =* 7.68 × 10^−137^). Similar patterns of upstream under-representation and downstream enrichment for guanosine have been reported in the literature (Bazak et al., 2014; Porath et al., 2014; Ramaswami et al., 2013). No motifs were observed at positions more distant than one nucleotide (Figure 1), an observation reported previously (Peng et al., 2012).

**Table 4.** Upstream and downstream nucleotide frequencies.

|  | A | C | G | U | Total |
| --- | --- | --- | --- | --- | --- |
| Upstream |  |  |  |  |  |
| All sites | 686 (34.3%) | 605 (30.2%) | 116 (5.8%) | 594 (29.7%) | 2,001 |
| A-to-I sites | 675 (34.6%) | 583 (29.9%) | 105 (5.4%) | 586 (30.1%) | 1,949 |
| Downstream |  |  |  |  |  |
| All sites | 351 (17.5%) | 317 (15.8%) | 987 (49.3%) | 346 (17.3%) | 2,001 |
| A-to-I sites | 344 (17.7%) | 301 (15.4%) | 970 (49.8%) | 334 (17.1%) | 1,949 |

The distributions of nucleotides immediately upstream (top) and downstream (bottom) of edited sites, considering both the set of all sites, and restricted to only A-to-I sites.

**Figure 1.**
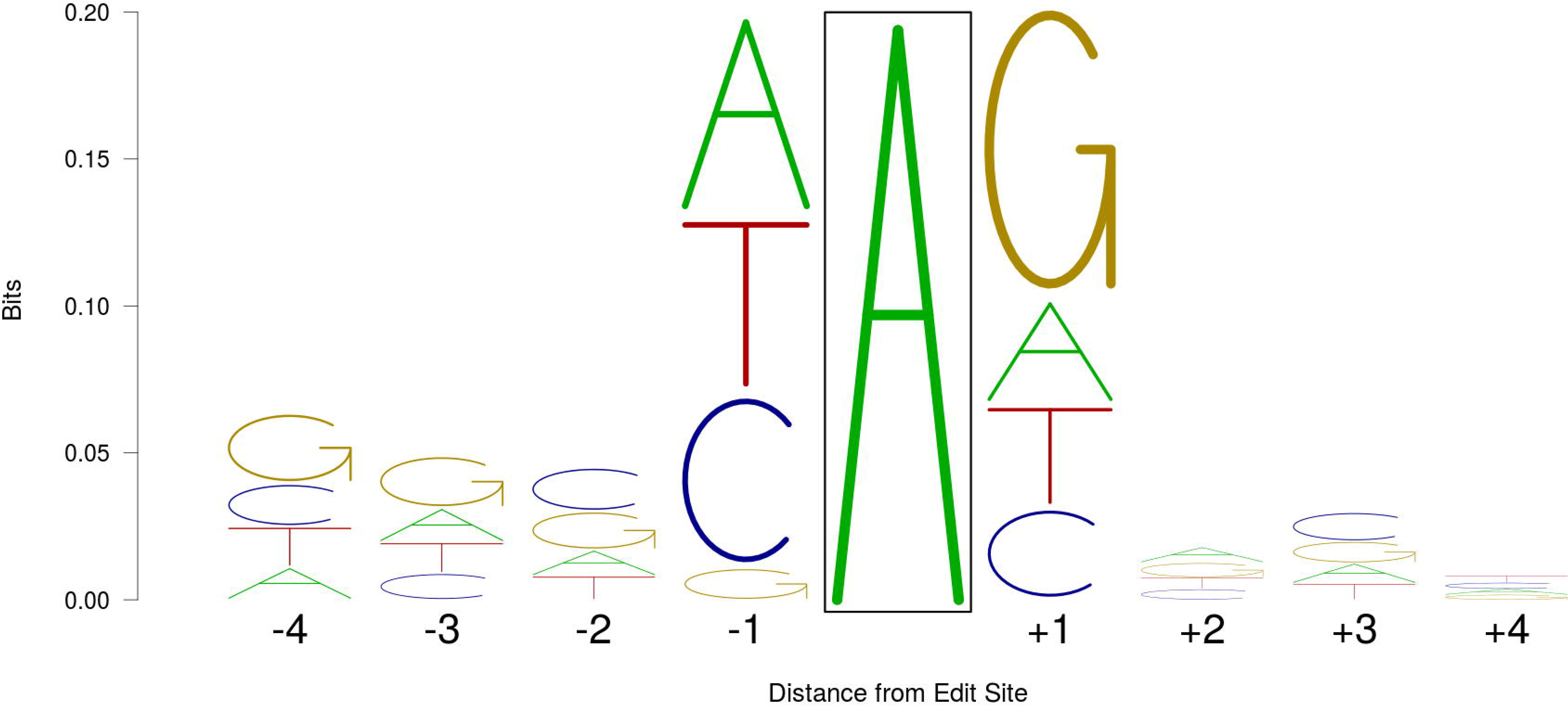
A sequence logo (Schneider and Stephens, 1990) showing the edit-containing consensus sequence based on all 1,947 A-to-I RNA edit sites identified in the current study. The proportion of each column occupied by each letter represents the frequency of that base at that position, while the total height of each column is equal to the information theoretical entropy (bits) at each position, with calculations as previously described (Schneider et al., 1986). For clarity of presentation, the edit site (boxed) is not shown at its actual height of 1.999 bits.

### Predicted edit sites occur predominantly in double-stranded regions

Since the ADAR1 and ADAR2 enzymes target double-stranded sections of RNA (Lehmann and Bass, 1999, 2000) it is expected that most edited sites will map to sequences able to form double-stranded structures. To test this hypothesis, double stranded regions of pre-mRNA transcripts were computationally predicted using R (R Core Team, 2017) to produce dot-plots. Figure 2 shows an example structure for *ABCG2*, a gene that encodes a transporter protein important to lactation and milk production ((Cohen-Zinder et al., 2005); plots for other structures are displayed in Figure S1). Visual examination of the predicted structures confirmed that the majority of candidate edited sites (85.1%; 1,703 of 2,001) are located within regions of RNA with the potential to form double-stranded helices. When only A-to-I edits were considered, an even higher percentage (87.1%; 1,698 of 1,949) were predicted to occur in such regions. Although some proportion of the 15% of sites not observed to reside in double stranded regions could be assumed to be false positive edit sites, these inconsistencies could arise from failure to accurately identify base-paired structures, or where the paired strand was more than 1,500 base pairs from any edit sites within the gene and therefore not included within the plotted region (see Materials and Methods for a description of double stranded prediction methodology). Images of the predicted double-stranded secondary structures are displayed in Figure S2.

**Figure 2.**
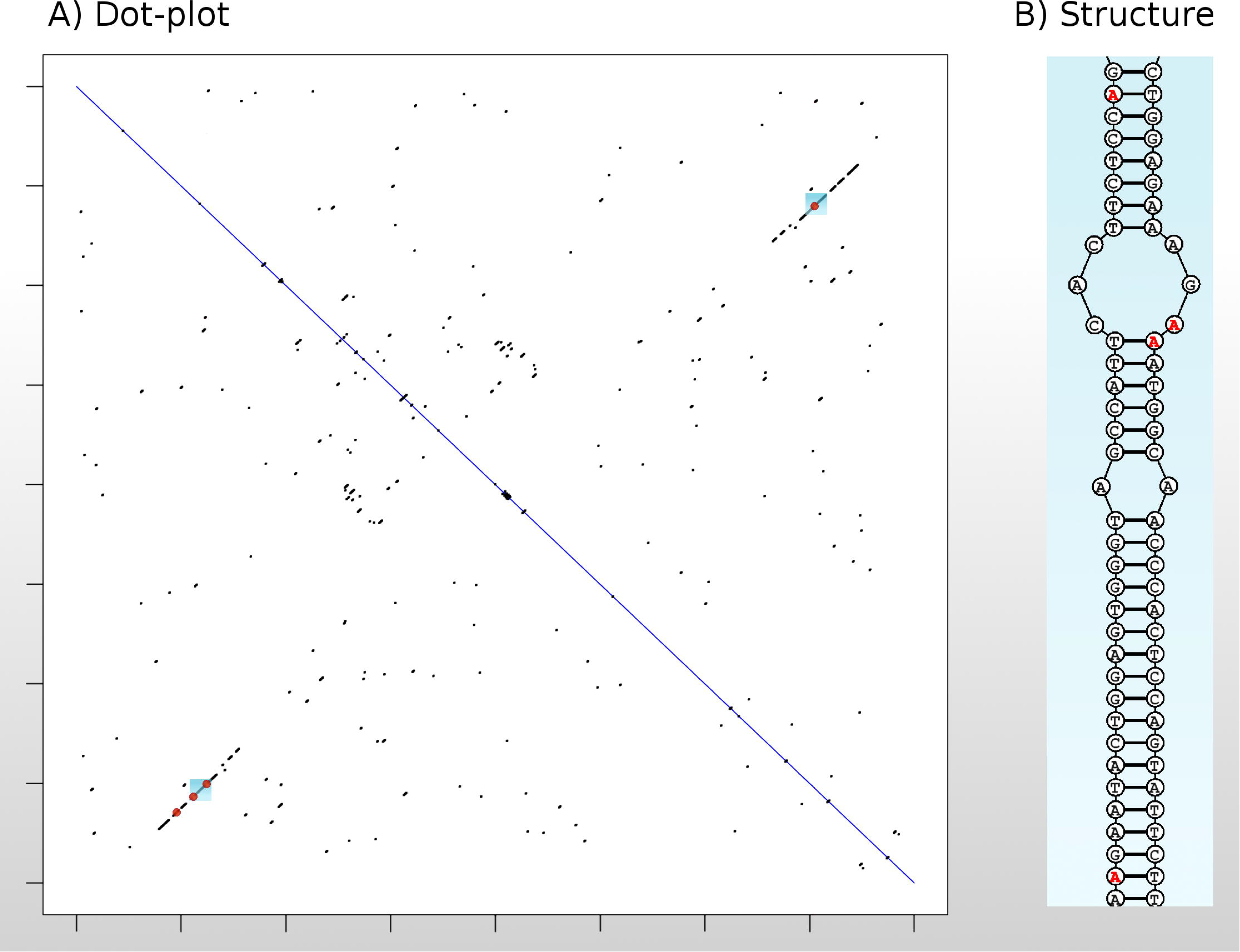
An example dot-plot showing visualisation of double-stranded regions of RNA sequences. A) A dot-plot for a 2 kbp section of the *ABCG2* gene, plotted against its complementary sequence. Black dots represent positions where at least eight of the surrounding eleven base pairs are complementary. Longer diagonal black lines indicate that long complementary sequences are present, representing helical regions in the secondary structure. Red dots highlight the positions of RNA edit sites. B) The secondary structure corresponding to the blue boxes highlighted in (A). Four RNA editing sites are indicated using red bases, corresponding to the red dots in (A). Structure (B) drawn using the ‘draw’ program from the RNAstructure (Reuter and Mathews, 2010) software package.

Within double-stranded regions, almost two-thirds of sites (n=1,127; 66.2%) were predicted to base-pair with a uridine residue, in accordance with standard Watson-Crick base-pairing rules (assuming an adenosine reference allele). The majority of the remaining sites were situated opposite to a cytidine residue (n=504; 29.7%), that would allow wobble base-pairing between the cytidine and inosine nucleotides after editing. The non A-to-I edit sites were much more sparsely represented within double-stranded regions, with only 9.6% (5 of 52) of the sites situated within these regions. This suggests that either the non A-to-I sites are edited by mechanisms which do not require double-stranded RNA, and/or that these sites have a considerably higher false positive rate than the A-to-I edits.

### Proportions of Reads Edited

To provide a quantitative assessment of editing in a larger population of animals, the base composition of candidate sites identified in the nine ‘discovery’ animals was assessed in 355 additional animals for which RNAseq data was available. The proportion of reads edited in these ‘quantification’ animals was defined as phi (Φ; (Park et al., 2017)). Phi values varied widely across sites: from 0.03% to 90.04% for A-to-I reads (median 12.81, mean 17.89). A significant association was observed between the upstream nucleotide at an editing site and the proportion of reads edited (*p* = 3.94 × 10^−13^, see Figure 3A). This was due primarily to an increase in editing where the upstream nucleotide is uridine (mean=21.72%), with a decrease observed for upstream guanosine (mean=13.33%). In contrast, no association was observed for the downstream adjacent nucleotide (p = 0.291, Figure 3B). Considering both the upstream and downstream bases simultaneously (Figure 3C), the highest rates of editing were observed at U.G and U.U sites, and the lowest at G.G and G.C sites, where the dot represents the editing site. Mean and median Φ values are displayed in Table S1 for all A-to-I edit sites.

Within double-stranded regions, lower values of Φ were observed (diff = −0.09, *p =* 0.018) where the predicted base pair of the edit site was a guanosine compared to other bases. These edit sites are also the least frequently observed, with only 15 observations among the 1,703 within double-stranded regions. Conversely, the most stable modification is expected when the paired base is a cytosine, and a significantly higher average Φ value was observed for these sites (diff = +0.05, *p =* 4.77 × 10^−10^).

**Figure 3.**
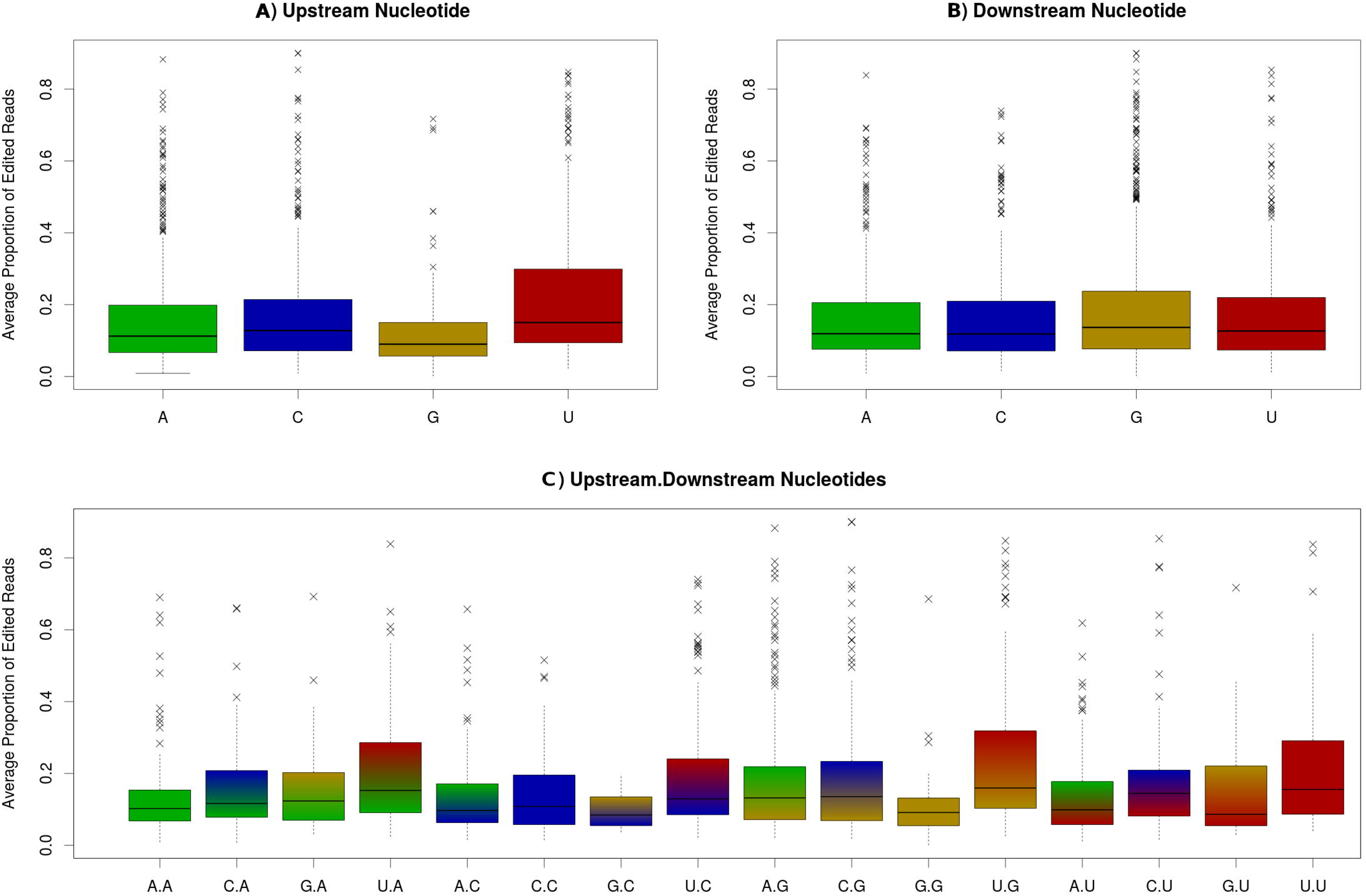
Distributions of the proportion of edited reads (Φ) for A-to-I RNA editing sites, by A) upstream nucleotide; B) downstream nucleotide; C) both upstream and downstream nucleotides. Proportions at each site are averaged across 355 animals in the quantification data set.

### Relationship between RNA-editing and transcript abundance

Given the observation of widespread editing across diverse mammary transcripts, we wondered what physiological effects might be attributable to these modifications. Since editing has previously been proposed as a mechanism to modulate gene expression through microRNA-based mechanisms (Liang and Landweber, 2007; Wang et al., 2013; Brümmer et al., 2017), or through nuclear-retention (Zhang and Carmichael, 2001; Prasanth et al., 2005), we looked at the relationship between Φ-values and transcript abundance by calculating Pearson correlation coefficients for each implicated gene. For this analysis, we were particularly interested in the impacts on mRNA, so to best represent spliced transcripts, transcript abundance was quantified using reads that either mapped wholly within exons, or mapped across exon-exon boundaries (see Methods and Materials). Strikingly, we noted significant correlations for a large proportion of edited transcripts (N=177 after Bonferroni adjustment; Figure 4), with the distribution of effects showing a strong bias towards genes whose expression was negatively correlated with Φ (Figure 4). Although it is unknown whether editing is driving these effects, these observations highlight a potential mechanism by which RNA editing may be impacting lactation phenotypes through modulation of mRNA abundance.

**Figure 4.**
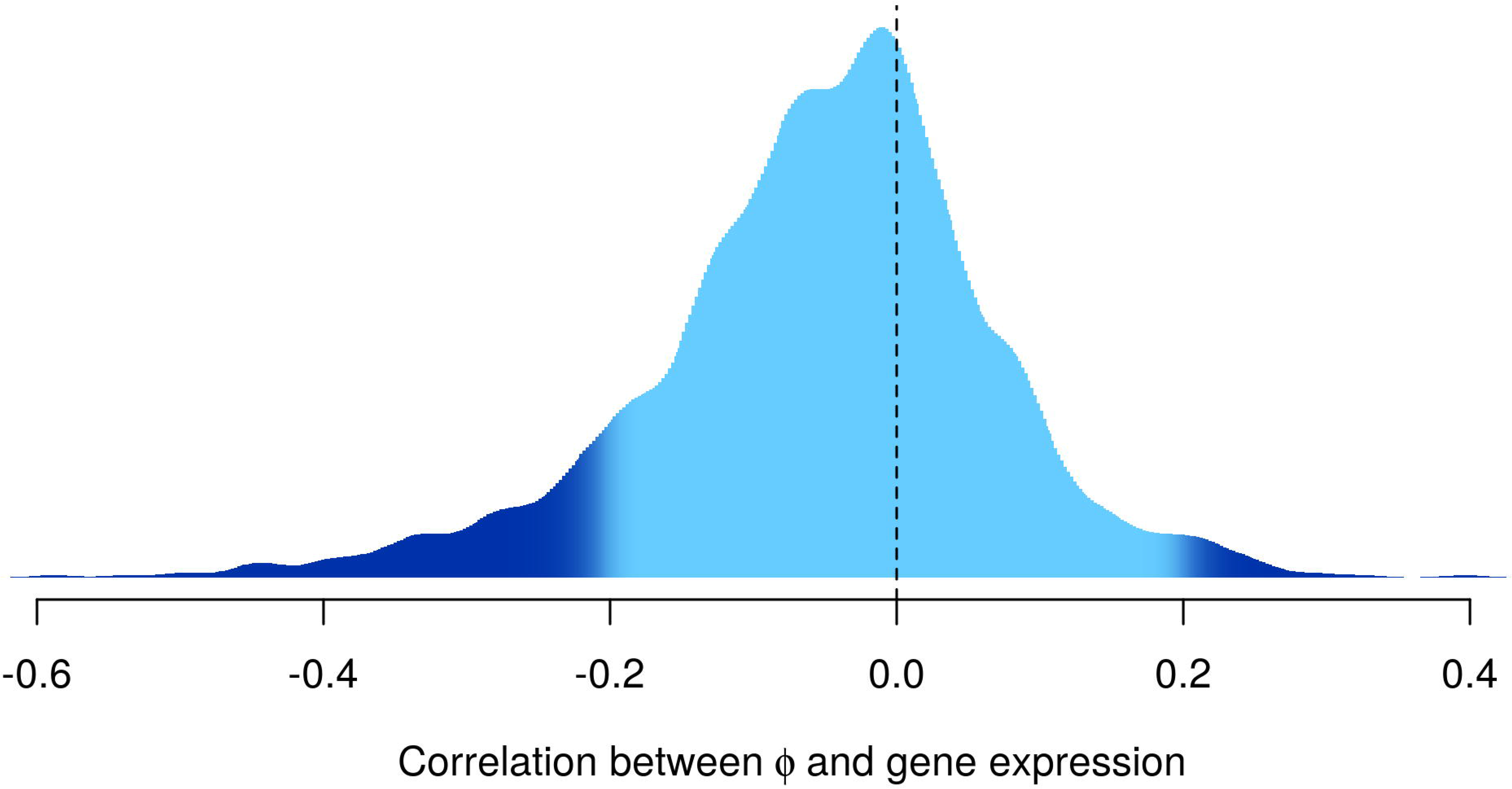
The distribution of Pearson correlation statistics calculated between editing rates (logit-transformed Φ values) and the expression of the genes to which they map (VST-transformed; see Materials and Methods). Dark blue indicates correlations which are significant after Bonferroni correction (P<2.62 × 10^−5^).

### Genome-wide association analysis of RNA edits

Having defined Φ values for all animals and all curated sites, we next used these data as phenotypes for genome-wide association studies (GWAS), with the aim of discovering RNA editing QTL. These models comprised generalised least squares (GLS) models, modelling the covariance between animals using a numerator relationship matrix based on pedigree records to account for underlying population structure and relatedness between animals (Materials and Methods). Using 630,774 genotypes from the Illumina BovineHD marker panel and logit-transformed Φ-values as phenotypes, 186 of the 2,001 RNA editing sites exhibited edQTL that were significant after Bonferroni adjustment for multiple testing (threshold 0.05/630, 774 = 7.93 × 10^−8^). Of these 186 edQTL, 131 sites harboured the top associated variant within 500 kbp of the editing site, and could therefore be assumed to be regulated in *cis.* These sites mapped to a total of 67 genes, with the *CSN3, ELF5*, and *PFKFB2*, each containing five or more associated sites. The full list of 131 sites is detailed in Table S2. Low levels of inflation in the test statistics were observed across the 2,001 edQTL GWAS (mean = 1.03, median = 1.02; ideal value 1.0), indicating that the generalised least squares models were adequately controlling for relatedness between the animals.

We next aimed to fine-map signals using imputed whole-genome sequence (WGS) data. Imputation was conducted using methods analogous to those previously described ((Lopdell et al., 2017); Materials and Methods). The 131 sites with *cis*-edQTL were remapped at WGS resolution for 1 Mbp windows in the 355 quantification set animals, with intervals centred on the most significant marker identified on the BovineHD panel for each site. Associations were conducted as per the analysis using the BovineHD panel. Thirty of 131 sites had at least one strongly associated WGS marker (exceeding the Bonferroni threshold) that mapped within the double-stranded region containing that site. Figure 5 shows example plots for the *HOOK3* gene.

**Figure 5.**
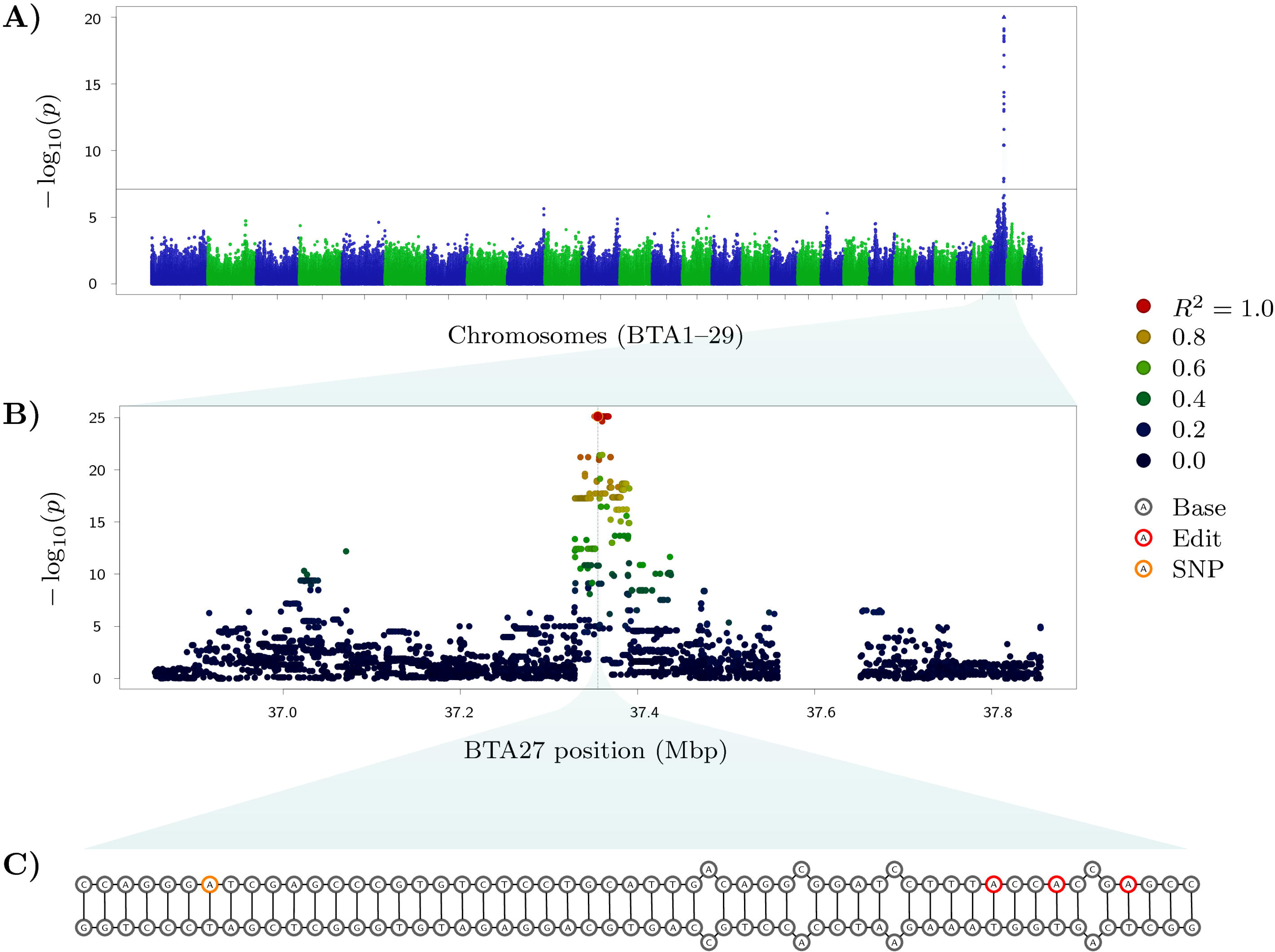
Genetic analysis of RNA editing at the *HOOK3* locus. A) A Manhattan plot showing an edQTL using markers from the BovineHD panel, with the edited site located at BTA27:37,355,505 within *HOOK3.* The horizontal black line indicates the genome-wide significance after Bonferroni correction. B) A 1 Mbp window centred on the most significant variant in the WGS-resolution data (rs109157662; BTA27:37,355,466). Colours represent the strength of LD (R^2^) with that variant. The vertical dashed line indicates the position of the edited site. No variants are present around 37.6 Mbp due to the presence of numerous small contigs at this locus in the reference sequence, which have been filtered out of the WGS data set due to errors in phasing. C) Putative structure of the pre-mRNA surrounding the edited sites (red) and candidate causative SNP (orange). Site BTA27:37,355,505 is the left-most edited site. The candidate causative SNP is also shown in (B) with an orange border.

### Examination of RNA phase and complementarity relationships between edit sites and candidate modulatory variants

Since base substitutions within double stranded RNA transcripts could be assumed to modify the structure and stability of such molecules, we reasoned that collocated, RNA-editing-associated WGS variants would make strong candidate causal variants for the observed edQTL. To investigate these relationships, edit sites that exhibited significant *cis*-edQTL were further analysed in the following two ways. First, read pair information was used to derive individual transcript haplotypes between edited bases and candidate causative alleles, with consistency of these phase relationships then assessed for heterozygous animals. Although we assumed such relationships would be due to impacts on base complementarity in double stranded RNA molecules, phase analysis was not restricted to regions predicted to form these structures, since the distance between edited bases and WGS variants was relatively short, and necessarily limited to the read fragment length (median unspliced length = 150 bp). Variant pairs were also filtered to exclude those that had fewer than five reads encompassing both sites. This yielded 48 pairs of edited bases and WGS variants, representing 26 distinct edit sites in 15 genes (where the reduced number of edits compared to pairs reflected sites that were paired with multiple variants). Association analysis revealed strong phase enrichment for the majority of pairs, with 43 of 48 significant at the Bonferroni threshold of P<0.001042 (see Materials and Methods).

The second analysis focused only on pairs of edited bases and edit-associated WGS variants collocating to double stranded structures (N=127 pairs; where double-stranded regions were predicted as previously described). We hypothesised that WGS alleles that were complementary to the base on the opposite, paired strand would increase the substrate affinity for ADAR enzymes, thus leading to increased editing for these sites (Figure 6). To test this, we removed all editing-associated variants for which neither allele paired with the opposite base on the complementary strand (N=109 pairs remaining). Using a one-sided t-test to assess whether the anticipated sign of effect between edits and complementary and non-complementary alleles was different than zero, a modest, but significant effect was observed (P=0.047). This observation, and the allele-specific editing results described above, supports the hypothesis that the mechanism of genetic modulation of editing at least partly derives from the impact of these variants on RNA secondary structures.

**Figure 6.**
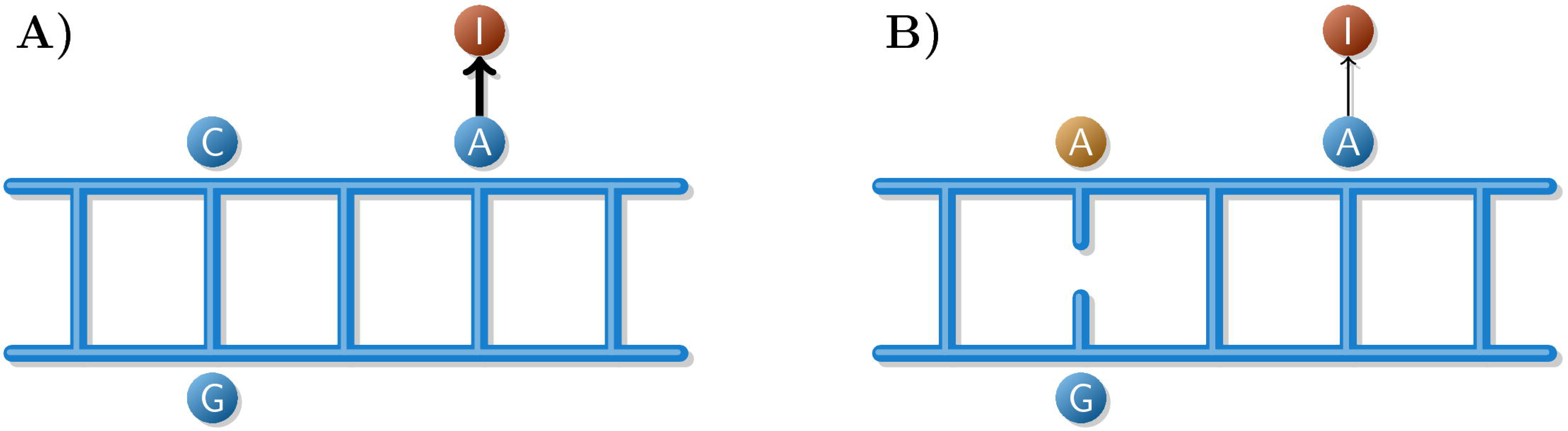
Graphical illustration of the mechanistic hypothesis that reducing sequence complementarity will decrease rates of editing. A) A short double-stranded section of an mRNA secondary structure, containing a SNP (C) with complementary (G), and an adjacent editing site (A), with high levels of editing. B) The alternative SNP allele (A) reduces complementarity, destabilising the secondary structure with a consequent reduction in editing rates.

### Correlations with expression and lactation QTL

RNA editing has previously been reported (Goldstein et al., 2017) to regulate levels of gene expression, so we hypothesised that edQTL may also influence expression, where these relationships should manifest as co-segregating edQTL and eQTL. To test this, we first analysed the 67 genes with *cis*-edQTL for the presence of *cis*-eQTL at WGS resolution. The methods used were analogous to those applied for detection of *cis*-edQTL, with gene expression phenotypes calculated from exonic read counts and normalised using the variance stabilising transformation in the DESeq R package (Anders and Huber, 2010, see Materials and Methods for further detail). This analysis revealed that 30 of the 67 genes had significant *cis*-eQTL with significance defined as having at least one variant with *p < 1 ×* 10^−8^. This list included genes with protein products known to be secreted in milk *(CSN3*: kappa-casein; *LPO:* lactoperoxidase; *LTF*: lactoferrin), along with several genes for which genetic impacts on milk composition or production have previously been published: *MARC1* (Lopdell et al., 2017), *SLC37A1* (Kemper et al., 2016), *STAT5B* (He et al., 2011).

To test for shared genetic architecture between the eQTL and collocated edQTL, Spearman correlations were calculated using the association χ^2^ statistics of each pair of QTL, in an approach similar to that previously described (Littlejohn et al., 2014b, 2016; Lopdell et al., 2017). This method highlights co-segregation patterns for QTL, where, assuming a common causal variant and haplotype structure between signals, the rank-order of associated markers should be similar for effects that are genetically co-regulated. Table S2 shows results for the 131 edits with *cis*-edQTL, alongside eQTL results and Spearman correlations between QTL pairs. At seven A-to-I edited sites (Table 5 and examples in Figure 7), correlations of greater than 0.7 were observed between edQTL and eQTL, potentially suggesting a gene expression consequence of the observed edQTL effects. These seven sites mapped to six discrete genes, with two sites impacting the *CSN3* gene.

**Figure 7.**
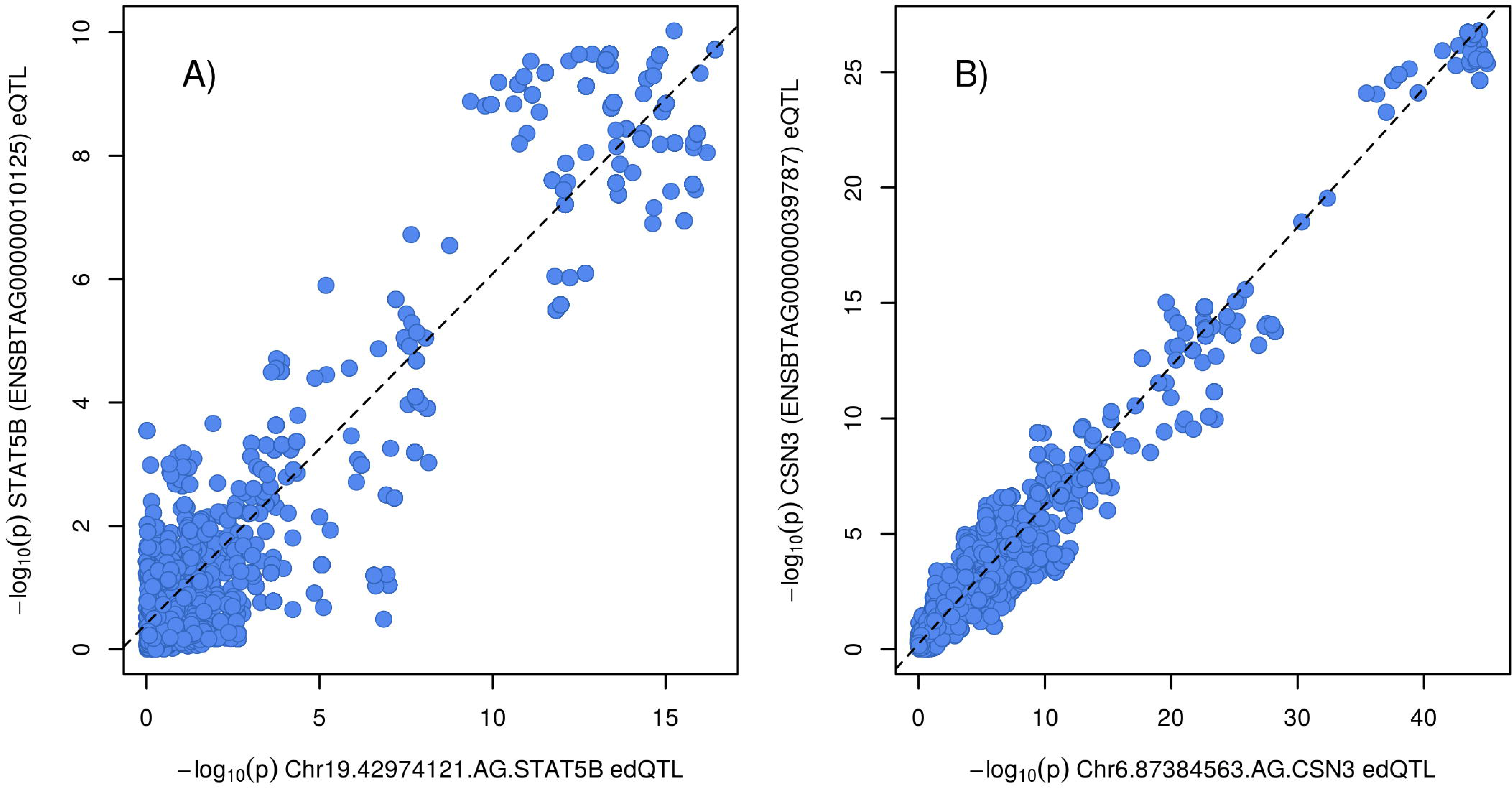
Two examples of co-located, co-segregating eQTL and edQTL. Each point represents the — log10 p-values for one variant for an edQTL (x-axis) and eQTL (y-axis). A) The edQTL for the Chr19.42974121.AG.STAT5B site, against the *STAT5B* eQTL, with correlation *r* = 0.814. B) The edQTL for the Chr6.87384563.AG.CSN3 site, against the *CSN3* gene, with correlation *r* = 0.906.

**Table 5.** Correlations between edQTL and eQTL.

| Site | Ensembl ID | Corr | eQTL Pval | edQTL Pval |
| --- | --- | --- | --- | --- |
| Chr6.87384501.AG.CSN3 | ENSBTAG00000039787 | 0.871 | $1.64 \times 10^{-27}$ | $2.31 \times 10^{-33}$ |
| Chr6.87384563.AG.CSN3 | ENSBTAG00000039787 | 0.906 | $1.64 \times 10^{-27}$ | $9.71 \times 10^{-46}$ |
| Chr6.99862424.AG.HPSE | ENSBTAG00000005745 | 0.915 | $2.65 \times 10^{-30}$ | $1.42 \times 10^{-38}$ |
| Chr16.4747421.AG.PFKFB2 | ENSBTAG00000002126 | 0.753 | $1.30 \times 10^{-9}$ | $3.96 \times 10^{-22}$ |
| Chr19.42974121.AG.STAT5B | ENSBTAG00000010125 | 0.814 | $9.44 \times 10^{-11}$ | $3.72 \times 10^{-17}$ |
| Chr19.9446655.AG.LPO | ENSBTAG00000012780 | 0.732 | $3.04 \times 10^{-18}$ | $1.02 \times 10^{-19}$ |
| Chr27.37355512.AG.HOOK3 | ENSBTAG00000007634 | 0.782 | $6.87 \times 10^{-20}$ | $7.98 \times 10^{-33}$ |

Spearman correlations between edQTL and eQTL for the gene in which the edited site was located. Results are shown only where sites i) were A-to-I edits; ii) were genome-wide significant for both the eQTL and edQTL; and iii) had correlations larger than 0.707 ( $r^2 > 0.5$ ).

Given the bias towards negatively correlated gene expression and editing *per se* (see ‘Relationship between RNA-editing and transcript abundance’ section above), we also wondered whether *cis*-edQTL/eQTL pairs would reflect this relationship, where we could anticipate allelic effects to show antagonistic signs of effect between edQTL and eQTL. To test this, pairs of eQTL and edQTL that were significantly correlated were identified (N=24; P<2.62 × 10^−5^) following Bonferroni adjustment), and subsequently classified as to whether the effect of the top edQTL variant had the same or opposite sign of effect to the eQTL. Given our prior hypothesis that the sign of effects would be reversed, we conducted a one-sided t-test that the mean sign of these effects was negative. This yielded a non-significant p-value of 0.114; however, repeating the analysis with a more conservative list of eQTL/edQTL pairs (100-fold smaller correlation p-value threshold of 2.62 × 10^−7^; N=15), yielded a highly significant P=0.007. This observation suggests that, at least for the loci for which eQTL/edQTL pairs are most strongly correlated (and thus most likely to represent a common genetic signal), increased levels of editing leads to decreases in mRNA expression.

Since edQTL might have further effects on lactation traits such as milk yield and milk component concentration phenotypes, we conducted association analysis on these phenotypes using imputed WGS genotypes and the GLS models described above. To perform association analysis of lactation traits, herd test data for 9,988 cows was used to test for the presence of fat, protein, lactose, and milk yield QTL that were collocated to each of the 131 *cis*-edQTL intervals. Examining co-segregation signals using the same methods applied to analysis of eQTL data, nine edited sites exhibited edQTL that were strongly correlated *(r* > 0.7) with at least one production QTL (Table 6 and examples in Figure 8). These effects were distributed across five genes, with three of these sites (in the *HPSE, STAT5B*, and *HOOK3* genes) also showing correlations with eQTL (compare Figure 8C and Figure 8D). Additional sites with correlations greater than 0.5 were observed in the *CSN3* and *LTF* genes (see Table S3).

**Table 6.** Correlations between edQTL and milk QTL.

| Site | MY | FY | PY | LY | FC | PC | LC |
| --- | --- | --- | --- | --- | --- | --- | --- |
| Chr4.40591602.AG.CD36 | **0.746** | -0.179 | 0.453 | 0.663 | **0.725** | 0.651 | 0.237 |
|  | 0.382 | -0.037 | 0.252 | 0.357 | 0.356 | 0.409 | 0.298 |
| Chr4.40592839.AG.CD36 | **0.753** | -0.210 | 0.458 | 0.678 | **0.733** | 0.644 | 0.223 |
|  | 0.362 | -0.051 | 0.286 | 0.323 | 0.339 | 0.236 | 0.353 |
| Chr6.99862424.AG.HPSE | 0.024 | 0.252 | -0.101 | 0.255 | -0.238 | **0.720** | 0.346 |
|  | -0.082 | 0.338 | -0.083 | 0.016 | -0.246 | 0.320 | 0.118 |
| Chr17.60341050.AG.FBXW8 | -0.294 | 0.237 | -0.336 | -0.323 | **0.753** | 0.592 | -0.216 |
|  | -0.305 | 0.092 | -0.442 | -0.356 | 0.258 | 0.479 | -0.082 |
| Chr17.60341051.AG.FBXW8 | -0.242 | 0.234 | -0.294 | -0.271 | **0.806** | 0.612 | -0.184 |
|  | -0.219 | 0.043 | -0.415 | -0.277 | 0.348 | 0.428 | -0.168 |
| Chr19.42974121.AG.STAT5B | -0.161 | -0.202 | -0.337 | 0.607 | -0.129 | **0.785** | **0.965** |
|  | 0.148 | -0.102 | -0.183 | 0.671 | 0.141 | **0.733** | **0.866** |
| Chr27.37355505.AG.HOOK3 | -0.166 | 0.639 | -0.153 | -0.142 | 0.395 | -0.172 | **0.709** |
|  | -0.163 | 0.351 | -0.117 | -0.151 | 0.121 | -0.021 | 0.419 |
| Chr27.37355508.AG.HOOK3 | -0.145 | **0.714** | -0.130 | -0.141 | 0.389 | -0.170 | 0.689 |
|  | -0.114 | 0.390 | -0.095 | -0.088 | 0.104 | -0.039 | 0.393 |
| Chr27.37355512.AG.HOOK3 | -0.180 | 0.668 | -0.171 | -0.156 | 0.417 | -0.170 | **0.738** |
|  | -0.179 | 0.377 | -0.179 | -0.147 | 0.219 | 0.020 | 0.472 |

Pearson (top row) and Spearman (bottom row) correlations ( $r$ ) between edQTL and milk production QTL. Only edited sites which i) had a significant edQTL; ii) were A-to-I edits; and iii) had at least one result where $r^2 > 0.5$ are shown. Results with $r^2 > 0.5$ are indicated in bold. Phenotypes are milk yield (MY), along with milkfat, protein, and lactose yield (FY, PY, LY) and concentration (FC, PC, LC).

**Figure 8.**
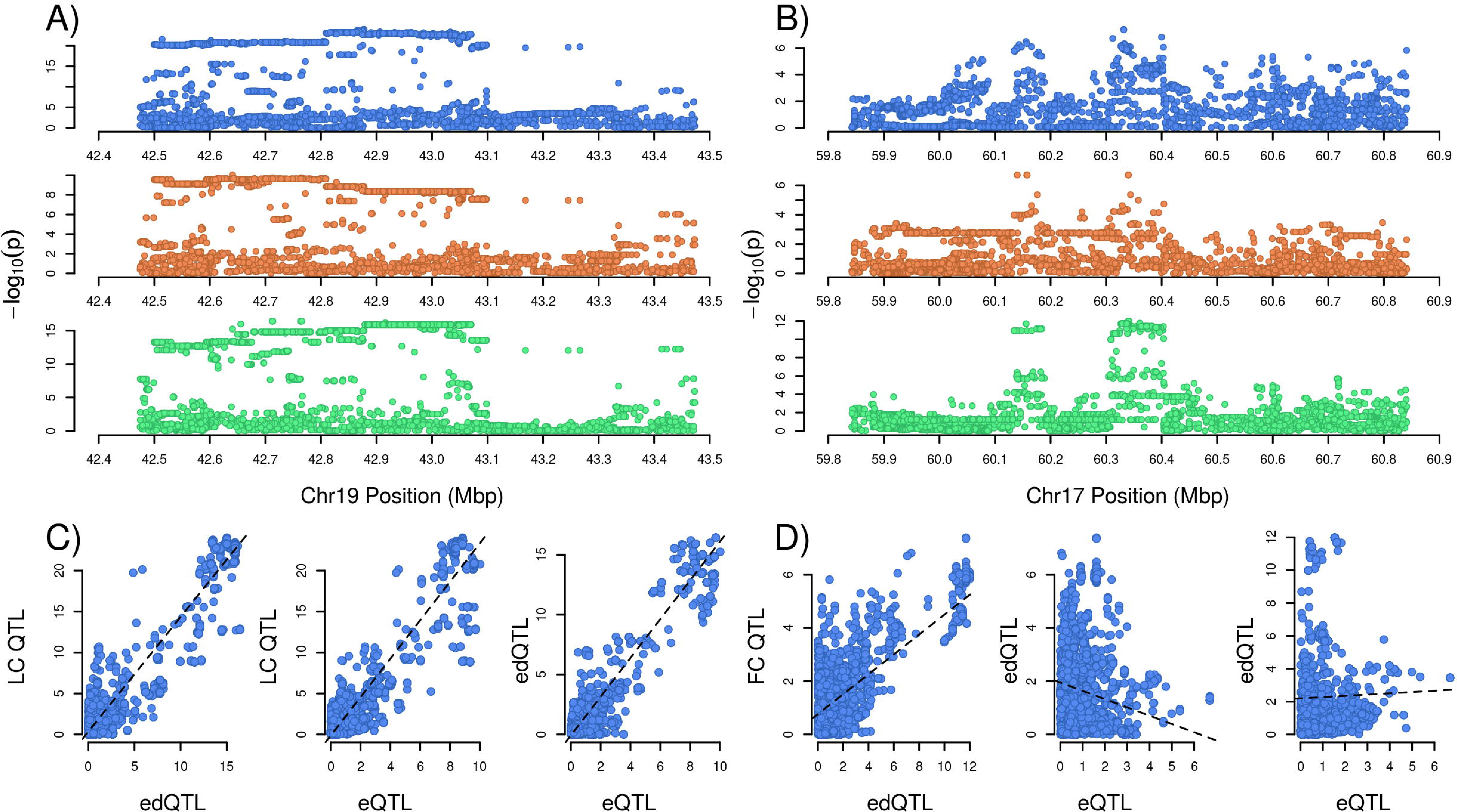
Production QTL and correlations with co-located edQTL and eQTL. Panels A and B: 1 Mbp windows for two production QTL (blue), eQTL (orange) and edQTL (green). Panel A shows the lactose concentration (LC) QTL at Chr19:42.9Mbp, with the co-segregating *STAT5B* eQTL and Chr19.42974121.AG.STAT5B site edQTL. Panel B shows the fat concentration (FC) QTL at Chr17:60.3Mbp, and co-segregating Chr17.60341051.AG.FBXW8 edQTL, along with an independently segregating *FBXW8* eQTL (Panel D). Panels C and D: plots of QTL p-values (— log_10_ scale) against one another. Panel C illustrates the strong correlations between all three QTL in panel A. Panel D shows a case where, while the FC QTL and Chr17.60341051.AG.FBXW8 edQTL are correlated, and therefore may share a similar genetic underpinning, the *FBXW8* eQTL is not correlated with either.

## Discussion

We report the discovery of 2,001 RNA editing sites in the bovine mammary transcriptome, and subsequently explore the genomic context and properties of these sites. We note strong correlations between the extent of RNA editing and the overall abundance of these transcripts, and we further report genome-wide association analyses of editing to identify genetic modulators of these effects. Association analysis of gene expression and lactation phenotypes for variants mapping to edQTL intervals reveals a number of overlapping signals at these locations, providing a potential mechanistic linkage between the editing of transcripts and mammary and lactation physiology. We discuss some of these findings in more detail, below.

### Editing site frequencies

The majority of editing sites discovered were A-to-I edits (1,941 of 2,001). This percentage (97.4%) is higher than the 80% reported previously for the cattle transcriptome (Chen et al., 2016), but is similar to results reported in human transcriptomes (97.25% (Porath et al., 2014); 93.8-99.2% (Chen, 2013)). In the cattle study referenced above (Chen et al., 2016), several tissues were examined, identifying between 180 and 404 edits per tissue. Studies conducted on human samples, by contrast, report far higher numbers of edit sites, where the numbers have increased over time due to the growing availability of large data sets: 14,500 in 2004 (Athanasiadis et al., 2004), 22,700 in 2012 (Peng et al., 2012), to over 100 million in 2013 (Bazak et al., 2014). The bulk of these edits occur in *Alu* repeat elements, where almost all adenosines are edited (Bazak et al., 2014). However, because these elements are primate-specific (Bazak et al., 2014), the observation of far fewer edits can be anticipated for cattle.

Previous work in mice (Gu et al., 2016) has reported the majority of edited sites falling in UTR regions. Here, in contrast, we found that the majority of sites (1,561 of 2,001) were intronic. This contrast may be partially attributable to differences between tissues and bovine and murine transcriptomes, though one important distinction between our study and that of Gu et al. (2016) is that we targeted much higher read depths (>200M reads per sample, versus 10M). The extreme read depth targeted in this study reflects a strategy to overcome expression biases that are a feature of secretory organs such as the lactating mammary gland (that present a limited diversity of highly expressed transcripts). It is fair to assume, however, that despite these biases, the increased read depth better represents intronic sequences in the current study.

The number of edited sites identified was also likely affected by the variant caller used to identify these sites, given the impact of allelic balance on calling variants. Edited sites, while resembling heterozygous SNPs, will in general not approximate the 50% allelic balance expected for variants in a diploid genome, with the balance dependent instead on the proportion of edited reads (Φ). Variant callers which are sensitive to allelic balance, including HaplotypeCaller (GATK RNAseq Best Practices), may fail to identify sites with low Φ values, so the results reported here likely under-represent sites with low levels of editing.

### Non A-to-I edits

In total, 52 (2.6%) of the 2,001 edited sites reported in this study were not canonical A-to-I edits. The three most common non-canonical edit types previously reported for cattle are C-to-U, G-to-A, and U-to-C (Chen et al., 2016), while G-to-A is also reported as the most common in humans (Porath et al., 2014). In the data reported here, the most common non-canonical edit type is also G-to-A. Functional edit sites of this type have been reported for humans in the *hnRNPK*, (Klimek-Tomczak et al., 2006), *TPH2* (Grohmann et al., 2010) and *WT1* (Niavarani et al., 2015) genes. These sites are hypothesised to be edited by the APOBEC3A enzyme in humans (Niavarani et al., 2015); however, the homologous cattle gene showed little or no expression in the mammary gland in the current study.

Twelve non-canonical edit sites exhibited C-to-U edits. This type of edit has been attributed in humans to the actions of the APOBEC1 enzyme (Siriwardena et al., 2016). The homologous bovine gene also shows minimal levels of transcription in the current study. Previous work has reported an over-representation of A and U nucleotides in the immediate vicinity of C-to-U edit sites in mice (Rosenberg et al., 2011), however, this was not observed for the sites detected in this study. Considering the minimal expression levels of APOBEC1, these observations suggest this class of edits may be enriched for false positive sites, and interpretation of these results should be considered accordingly in our analysis.

### Relationships between editing, double-strandedness, and transcript abundance

The majority of edited sites were located within regions for which double-stranded secondary structures were predicted. Of these sites, over 66% were adenosine nucleotides base-paired to uridine nucleotides, in accordance with Watson-Crick pairing rules. When these sites are edited, the stability of the resulting structure is likely to be reduced, though it is noteworthy that I-U base pairs are valid under wobble base-pairing rules (Murphy and Ramakrishnan, 2004). Conversely, almost 30% of sites were predicted A-C pairs, which we expect to be unstable until edited into the I-C wobble base-pair. Therefore, we hypothesise that RNA editing is contributing to modulation of the stability of folded pre-mRNA secondary structures. Lower editing frequencies (Φ) were observed in double-stranded regions where the predicted base pair of the edit site was a guanosine. This observation can be explained by wobble base pairing (Murphy and Ramakrishnan, 2004), as guanosine is the only standard RNA base which does not pair with inosine, resulting in lower stability in the double-stranded region after editing at these sites compared to sites paired with other nucleotides.

We also noted that for many genes, the proportion of editing was significantly correlated with transcript abundance. These correlations were largely negative, where increased editing was associated with decreased mRNA expression. A biological cause and effect relationship is difficult to establish here, given that RNA editing may be a consequence (as opposed to cause) of reduced mRNA production, and other biases relating to library preparation or sequencing (Harcourt et al., 2017) could conceivably lead to spurious, negative correlation. However, these findings are similar to those reported in other analyses of global RNA editing profiles (Hwang et al., 2016), where a broadly negative relationship between editing and transcript abundance also fits with a mechanism by which mRNA expression is controlled through preferential retention of edited transcripts in the nucleus (Zhang and Carmichael, 2001; Prasanth et al., 2005).

### RNA edited genes and QTL

A number of the editing sites we detected mapped to genes that are involved in lactation. These genes include the major milk protein components (caseins) encoded by the *CSN1S1, CSN2*, and *CSN3* genes, as well as the antimicrobial LPO and LTF proteins. Edited sites were also observed in *STAT5A* and *STAT5B*, comprising transcription factors with critical roles in mammary differentiation and lactation (Liu et al., 1995; Cui et al., 2004). A number of other edited genes are important mediators of milk fat synthesis (*LPL*, *ACACA, GPAM* (Bionaz and Loor, 2008)) and secretion *(XDH, PLIN3* (Bionaz and Loor, 2008)), or are involved in the transport of small molecules in milk: *SLC37A1* (Pan et al., 2011), *ABCG2* (Otero et al., 2016). Together, these genes represent some of the most prominent and well-published genes in lactation biology, and include many of the largest effect loci implicated in genetic regulation of these traits (He et al., 2011; Khatib et al., 2008; Bionaz and Loor, 2008; Kemper et al., 2016; Olsen et al., 2007). This suggests, at a minimum, that RNA editing may functionally moderate aspects of mammary and lactation physiology, and further presents RNA editing as one of the mechanisms that may underpin milk and lactation QTL.

To investigate this hypothesis, GWAS was conducted for all 2,001 edited sites, treating the RNA editing proportion (Φ) as a phenotype. This analysis yielded significant *cis-* edQTL at 131 sites. Further analysis of these edQTL suggested that highly associated variants tend to be in phase with the corresponding edit sites, implying a consistency of phase within individual pre-mRNA molecules. We also found evidence that, when edit sites are predicted to be located within the same double-stranded secondary structure as a significant variant, alleles which increase the stability of the structure tend to increase editing, and alleles which destabilise the structure tend to decrease editing. These results are concordant with a mechanism whereby *cis*-edQTL causal variants act within each pre-mRNA transcript to stabilise or destabilise secondary structures, potentially modifying the substrate affinity of these molecules to RNA editing enzymes. These findings broadly support an analysis of the genetic impacts of RNA editing in humans, where the authors similarly looked at aspects of allele-specific editing (Park et al., 2017). This human study (Park et al., 2017), and two other studies in mouse (Gu et al., 2016) and *Drosophila* (Ramaswami et al., 2015), similarly propose allelic effects on folded RNA structures as the likely mechanism driving edQTL.

To look for potential impacts of edQTL on gene expression and lactation phenotypes, we conducted association mapping using intervals of WGS-resolution variants highlighted from RNA-editing analyses. Of the 67 genes highlighted with *cis*-edQTL, we identified 30 with significant *cis*-eQTL, seven of which also showed strong correlation of association statistics (*r* > 0.7). We also assessed correlations between edQTL and collocated lactation QTL, determined in a separate population of 9,988 outbred cows. Correlations greater than 0.7 were observed for A-to-I sites in the *CD36, FBXW8, HOOK3, HPSE*, and *STAT5B* genes. Two sites in the *CD36* gene were correlated with collocated fat concentration and milk yield QTL, with lower correlations also observed for the lactose yield and protein concentration phenotypes. This gene encodes a glycoprotein that has been implicated in fatty acid transport (Ibrahimi and Abumrad, 2002), and in mammary gland cell proliferation and involution (Spitsberg et al., 1995). The *HOOK3* gene contained three sites with edQTL that were strongly correlated with QTL for lactose concentration or fat yield. *HOOK3* encodes a microtubule tethering protein involved in intracellular vesicle trafficking (Xu et al., 2008), and is broadly analogous in function to the gene *PICALM* that has previously been associated with lactose concentration in milk (Lopdell et al., 2017). Of the five genes with strong correlations with lactation

QTL, *CD36* and *FBXW8* had relatively modest edQTL/eQTL correlations, whereas *HPSE, HOOK3*, and *STAT5B* showed correspondingly strong correlations with eQTL. For these latter genes, our findings present a potential chain of causality from variants modulating the editing of pre-mRNA transcripts, to consequent mRNA expression and lactation effects. Although the mechanism linking mRNA expression to physiological impacts is straightforward, the understanding of the impacts of RNA editing on gene expression is comparatively poor, though can be expected to advance in accordance with the rapidly growing body of literature regarding RNA editing biology. Together, these results improve our understanding of RNA editing in mammals, and our understanding of the link between genotypes and phenotypes in lactation.

## Materials and Methods

### DNA and RNA sequencing

Potential RNA editing sites were detected by comparing variant calls from mammary RNAseq to whole-genome DNA sequence calls. A total of 364 cows of mixed Holstein-Friesian, Jersey and cross-bred ancestry were divided into two non-overlapping sets. A discovery set composed of nine F2 Holstein-Friesian/Jersey cross-bred animals was sequenced using both RNAseq and whole-genome approaches, to enable discovery of edited sites within the RNA. The second set of 355 animals (the quantification set) was sequenced using RNAseq only, and used to quantify the level of editing at the sites first identified in the discovery set. The quantification animals were used to generate editing proportion phenotypes (Φ) for use in edQTL mapping.

RNA sequencing was performed on mammary biopsies from all 364 animals, as reported previously (Littlejohn et al., 2016). Briefly, high-depth mammary RNAseq was conducted on tissue obtained via mammary biopsy, sampled at several points in time. Following library preparation, samples were sequenced using the Illumina HiSeq 2000 instrument to produce 100 bp paired-end reads. RNASeq reads were mapped to the UMD 3.1 reference genome using Tophat2 (version 2.0.12) (Kim et al., 2013), mapping an average of 207.9 million reads per sample. Duplicate reads were marked using the MarkDuplicates command in the Picard software package (version 1.89; Broad Institute).

Whole genome sequencing was performed for the animals in the discovery set using methods we have described previously (Littlejohn et al., 2014b, 2016). All animals were sequenced using 100 bp paired-end reads on the Illumina HiSeq 2000 instrument, followed by mapping to the UMD 3.1 bovine reference, using BWA MEM 0.7.8 (Li and Durbin, 2009). This yielded mean and median mapped read depths of 22.1× and 22.2× respectively.

### Identifying edited sites in the RNA

Variant calling was performed on the discovery set animals, for both the DNA and RNA alignments, using GATK HaplotypeCaller version 3.2 (DePristo et al., 2011). Reads that had been marked as duplicates, or with mapping quality scores below twenty, were excluded. Variants present in the dbSNP database (build 146) were also excluded. Additional filters were subsequently applied to the RNAseq variant calls, excluding variants with quality scores less than 100, cumulative read depth less than 135 (average 15 reads per animal), and any variants with five or fewer observed alternative alleles. Less stringent filters were applied to the WGS-called variants, excluding those with quality scores less than 50 or cumulative read depths less than fifty. Due to the difficulty in accurately calling indel variants, these were excluded from both variant sets.

Variants present in the RNA-called set but absent from the DNA-called set formed the initial set of potential RNA edits. This set was filtered further by removing any variants which had been called from WGS in a separate larger study (Littlejohn et al., 2016), yielding a set of 3,280 candidate RNA editing sites. These sites were further subjected to manual evaluation to remove sites where, for example, no read coverage was available in the DNA sequence, or where alignments in the RNA sequence appeared anomalous. This resulted in a conservative subset of 1,171 sites. During this process, we also observed numerous additional sites present in close proximity to manually inspected sites. These bases had not been annotated by the variant caller, likely as a consequence of under representation of the non-reference ‘allele’ (mean Φ = 0.22 and 0.17 for GATK-annotated and un-annotated sites respectively). These sites were also added to the curated set, providing that at least one edited read was present at the site in at least five of the nine discovery animals. Using these criteria yielded a final set of 2,001 manually appraised sites. Due to the conservative process for identifying and manually curating sites, and the subsequent incorporation of sites discovered by virtue of collocation with highly edited bases, our dataset is likely biased towards the most highly edited transcripts. Although use of an alternative variant caller that was less sensitive to allele biases may have enabled discovery of larger numbers of RNA edits genome-wide, this would have represented a compromise given the lack of unity between different variant callers, and consequent increased false positive rates. An example region comparing DNA and RNA sequencing data for three animals is illustrated in Figure S3.

Editing proportions for each of the 2,001 verified sites were calculated for each cow in the quantification set by reporting the base composition of reads in the RNA alignments. Edit sites were allocated to genes using the Ensembl Variant Effect Predictor (McLaren et al., 2016), requiring genes to map on the correct strand. Because UTR regions in the released annotations often appeared to be considerably shorter than those evident in the RNA sequence data, variants labelled as upstream or downstream were considered to sit in 5’ and 3’ UTRs respectively, given that they were discovered in RNA (i.e. expressed) data.

### Predicting two-dimensional mRNA structure

Local mRNA secondary folding structure was predicted for each editing site. Within each gene, sequence was extracted for an interval that included 1.5 kbp of sequence upstream and downstream of the most 5’ and 3’ edited sites. In cases where the total sequence extracted for a gene exceeded 15 kbp, multiple, shorter sub-sequences were used.

Each sequence was then plotted against its complement to generate dot-plots (Figure 2 and Figure S1). Dots were placed where at least 11 of the 15 nucleotides, centred on each pair of positions, were complementary. Diagonal lines appearing in the plots are indicative of long strands of complementary sequence, highlighting potential doublestranded regions for examination, by manual observation, for the presence of edited sites. These regions were also processed using the bifold-smp program from the RNAstructure software package (Reuter and Mathews, 2010) to generate candidate secondary folded structures.

### Genotyping and RNA QTL discovery

To enable the discovery of edQTL via GWAS, all 355 animals in the quantification data set were genotyped using the BovineHD SNP-chip (Illumina). Variants with minor allele frequencies < 1% were removed. As a filter for erroneous SNP assays, tests were conducted for Hardy-Weinberg equilibrium (Graffelman and Moreno, 2013) using PLINK software (Chang et al., 2015; Purcell et al., 2007) (version 1.9b3i), with variants yielding p-values less than 1.0 × 10^−30^ excluded. The final variant set, containing 630,774 variants, was used for edQTL and eQTL.

As described above, the base composition of RNA editing sites was determined in the quantification set of animals to determine the proportions of edited reads for each animal and site (Φ, (Park et al., 2017)). To satisfy the normality requirement for phenotypes used in the generalised least-squares model, the proportions of edited reads were transformed using the logit function. For each edited site, logit-transformed Φ values (y) were fitted to a generalised least-squares model to identify edQTL. The numerator relationship (A) matrix was used to account for any covariances between animals that were due to shared

Each genotyped variant was fitted individually using the generalised least-squares (GLS) model described in (Lopdell et al., 2017). Briefly, the model used was y = **X*β*** + *ϵ*, where the error term (*ϵ*) has the conditional mean 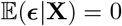 and covariance Var(*ϵ*|**X**) = **W**, where 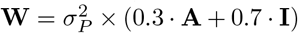, **I** is the identity matrix and 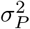 is the phenotypic variance.

To confirm that shared ancestry was not inflating the GLS results, the statistic 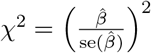 was calculated for each variant using the estimate of the slope 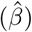. For each edited site, the median of the χ^2^ statistic was calculated, with the ratio of the observed median and the expected median (0.4549) yielding the inflation statistic for that edited site. Inflation is indicated when the value of this statistic exceeds unity.

As part of a previous study (Littlejohn et al., 2016), gene expression phenotypes were derived for a larger set of 375 animals, of which the 355 animals in the quantification set formed a subset. Gene expression, measured in fragments per kilobase of transcript per million mapped reads (FPKM), and in transcripts per million (TPM) (Wagner et al., 2012), was quantified for genes containing RNA editing sites using Stringtie software (version 1.2.4) (Pertea et al., 2015), with annotations from Ensembl genebuild release 81. Additional gene expression phenotypes were also derived by applying the “variance stabilising transformation” (VST) function from DESeq (Anders and Huber, 2010) to read counts for each gene, resulting in phenotypes with a distribution closer to the normal distribution, and therefore more suitable for analysis with linear models. The read counts used here consisted of only those reads that either a), mapped entirely within a single exon; or b), spliced across one or more annotated exon junctions, according to the exon boundaries defined by the Ensembl annotations (release 81). Reads that spliced at a site not recorded in the Ensembl annotations were excluded.

### WGS imputation and fine mapping

To map variants at a higher resolution around identified edQTL, WGS variants were imputed into the quantification animal set using a previously described reference population of 565 animals (Littlejohn et al., 2014a, 2016), comprising Holstein-Friesians, Jerseys, and cross-bred cattle. Briefly, these cattle were sequenced using the Illumina HiSeq 2000 instrument, yielding 100 bp paired-end reads. Mapping was conducted using BWA MEM 0.7.8 (Li and Durbin, 2009), resulting in mean and median mapped read depths of 15 × and 8× respectively. Variant calling was conducted using GATK HaplotypeCaller (version 3.2) (DePristo et al., 2011) with base quality score recalibration, followed by phasing using Beagle 4 (Browning and Browning, 2009). Variants with phasing allelic R^2^ < 0.95 were excluded for quality filtering purposes.

1 Mbp intervals, centred on the top *cis*-edQTL markers, were imputed to whole-genome sequence resolution using Beagle 4 (Browning and Browning, 2009) with the reference population described above, excluding variants with minor allele frequencies below 0.05. Across all 131 intervals, this process resulted in a total of 659,199 variants (mean 5,032; min 2,102; max 10,870 per interval). Although we have no truth set with which to directly determine the imputation accuracy for these animals, previous work we have performed (Littlejohn et al., 2016) indicates accuracies of around 98-99% when imputing BovineHD-resolution genotypes to WGS. Association analysis was conducted using the same GLS model described for SNP-chip based GWAS. Within the same intervals, gene expression phenotypes (described above) were used analogously to discover eQTL.

### Phase and complementarity relationships

To investigate phase relationships between edited sites and variants on the same transcript, WGS variants for each site with a significant edQTL were extracted. These analyses were restricted to animals heterozygous for the implicated site, and only variants that were correlated *R*^2^ > 0.95 with the most significant variant, and were within 150 bases of the edit site, were included in these analyses. These criteria yielded 59 pairs of edit sites and WGS variants. Read pairs were then extracted from RNAseq BAM files where the read pairs contained both the edit and variant positions. Reads meeting these criteria were subsequently counted to yield 2 × 2-contingency tables for the number edited / not edited for the edit site, and the number reference / alternative for the variant allele. Contingency tables of expected counts under independence were generated, and any pairs where at least one cell in either the observed or expected contingency table was less than six were excluded. This yielded 48 pairs that were then tested for independence using a t^2^ test. Results where *P* < 0.001 were considered significant, applying a multiple testing threshold of *P* = 0.05/48.

Complementarity relationships were investigated by extracting pairs of edit sites that exhibited significant edQTL, and WGS variants that collocated to the same doublestranded secondary structure. Structures were determined using dot-plots and the RNAstructure software package (Reuter and Mathews, 2010) as described above. Only WGS variants within 8 kbp of the edited site were considered. Additive allelic substitution effects (β) for the non-complementary allele were extracted from the WGS-resolution edQTL analysis for each edit site / variant pair. As we hypothesised that decreased complementarity would decrease editing, a one-sample í-test was performed for the one-sided alternative hypothesis that mean *β <* 0.

### Milk phenotypes and QTL

To examine the effect of RNA editing on milk production traits, milk fat, protein and lactose phenotypes were derived for 9,988 cows from measurements taken as part of standard herd-testing procedures. Milk samples were processed by LIC Testlink (Newstead, Hamilton, New Zealand) using Fourier transform infrared spectroscopy on Milkoscan FT6000 (FOSS, Hillerød, Denmark) and Bentley FTS (Bentley, Chaska, USA) instruments. Individual phenotypic measurements for each animal were estimated using repeated measures models in ASReml-R as described in (Lopdell et al., 2017).

These 9,988 cows had previously been genotyped on a mix of bovine SNP platforms: Illumina Bovine SNP50 (N=6,481), BovineHD (N=62), and GeneSeek Genomic Profiler BeadChip (N=3,949; GeneSeek/Illumina). Five hundred and one cows had been genotyped on multiple panels. All cows were imputed to WGS resolution for the 131 edQTL intervals using Beagle 4 as described above. These genotypes were subsequently used with the milk-sample-derived phenotypes to explore QTL at each of these intervals, using the GLS model described above.

## Acknowledgements

The authors would like to acknowledge S. Morgan and staff at DairyNZ Ltd. (Hamilton, New Zealand), and Phil McKinnon, Ali Cullum and staff at AgResearch (Hamilton, New Zealand) for facilitating mammary tissue sampling of lactating animals. We also wish to acknowledge New Zealand Genomics Limited (NZGL) and the University of Auckland Centre for Genomics, Proteomics, and Metabolomics for RNA preparation and sequencing, as well as both the Australian Genome Research Facility (AGRF) and Illumina FastTrack for both RNA and genomic DNA sequencing. This work was supported by the Ministry for Primary Industries (Wellington, New Zealand), who co-funded the work through the Primary Growth Partnership.

## Declarations

### List of abbreviations

edQTL: RNA editing quantitative trait locus; eQTL: gene expression quantitative trait locus; FPKM: fragments per kilobase of transcript per million mapped reads; GWAS: genome-wide association study; miRNA: micro-RNA; TPM: transcripts per million; UTR: untranslated region; WGS: whole genome sequence

### Ethics approval

All animal experiments in this study were conducted in accordance with all rules and guidelines in the New Zealand Animal Welfare Act 1999. As the majority of data were generated as part of routine commercial activities, formal committee assessment and ethical approval (as defined by the above guidelines) were not required. Samples were obtained for the mammary tissue biopsy experiment in accordance with protocols approved by the Ruakura Animal Ethics Committee, Hamilton, New Zealand (approval number AEC 12845). No animals were sacrificed for this study.

### Data Availability

Sequence (BAM) files containing the WGS and RNAseq reads sequenced from the nine discovery set cows, have been uploaded to the NCBI sequence read archive (SRA). BioProject accession number PRJNA446068 (SRP136662), BioSample accession numbers SAMN08810150 to SAMN08810167.

## Supporting Information

**Figure S1**

Dot plots of genome sequences centred on RNA editing sites. Sequences are plotted against their complement, with dots indicating that at least 11 of the 15 surrounding nucleotides are complementary, yielding diagonal black lines where long strands of complementarity are present, indicating potential double-stranded regions. Red dotted lines indicate the positions of RNA editing sites, showing that these tend to cluster within the double-stranded regions.

**Figure S2**

Structures predicted using the ‘bifold’ program from the RNAstructure (Reuter and Mathews, 2010) software package, for double-stranded regions predicted from the dot plots shown in Figure S1. Edited sites are indicated in red.

**Figure S3**

An example comparison between WGS and RNAseq for three animals, illustrating the difference between SNPs and RNA editing. The region shown is part of intron 1 of the LPO (lactoperoxidase) gene. The top row shows the single SNP called in this region from a large WGS study. The section with the blue background shows the WGS coverage mapped for three animals, where grey represents the reference base (indicated at the bottom of the figure), and with blue and brown representing cytosine and guanine respectively. The section with the yellow background shows the RNAseq coverage for the same three animals (green = adenine). Edit sites appear in the RNAseq coverage as bars of mixed green and brown (A-to-G(I) edits), while SNPs appear at the same location in both the WGS and RNAseq sequences.

**Table S1**

Summary data for edited sites. Tab one (“Edited Sites”) contains the chromosome and base position of each site on the bovine UMD 3.1 reference genome, along with the gene in which the site is located by HGNC symbol and Ensembl ID, plus the reference and edited base, complemented when the strand is negative. Tab two (“VEP Results”) contains the outputs from the Variant Effect Predictor for each edited site. The reference and edited base are not complemented in this tab. Tab three (“Edit Frequencies”) contains the data in tab one, with the addition of the immediate upstream and downstream bases, plus the mean, median and standard deviation of the editing frequency for each site. Only A-to-I sites are included.

**Table S2**

Details of the 131 edit sites exhibiting genome-wide significant *cis*-edQTL. Gene symbols and Ensembl identifications are provided for genes containing the edit sites. Also included are the minimum p-values for each edQTL, as well as the strength of the *cis*-eQTL for the appropriate gene. The Spearman correlation between the association (χ^2^) statistics for the edQTL and eQTL are also included, in the second tab.

**Table S3**

Correlations between all 131 *cis*-edQTL and milk production QTL. Edit sites are named by position (UMD 3.1 reference), reference and edited bases, and gene symbol. Tab one contains Pearson correlations, and tab two contains Spearman correlations. On both tabs, correlations greater than 0.707 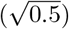 are highlighted in bold, those greater than 0.5 are italicised, and those less than zero are in grey.

